# Computational design of microbial and animal rhodopsin soluble analogues

**DOI:** 10.64898/2026.06.27.734960

**Authors:** Alexander T. Hilditch, Casper A. Goverde, Natalia Gasilova, Sarah Warrelmann, Nicolas Goldbach, Josef Wachtveitl, Laure Menin, Bruno E. Correia

## Abstract

The computational design of soluble analogues of membrane proteins has unlocked exciting opportunities for the integration of unique membrane protein functions into soluble proteins. Here, we use AF2_seq_ to generate accurate soluble analogues of both animal and microbial rhodopsins, based on the membrane GPCR topology, and the microbial rhodopsin transmembrane fold. We characterize the analogues and demonstrate that they are well-folded and highly thermostable. Furthermore, they exhibit the expected red shift characteristic of retinal binding. Top-down mass spectrometry confirms the placement of retinal covalent attachment, while X-ray crystallography validates the structural fidelity of the microbial rhodopsin analogue. Notably, the microbial rhodopsin analogue retains the primary reaction of the retinal photocycle, closely matching that of the native membrane protein. Overall, this work advances the possibility to transfer unique membrane protein functions, such as retinal photoswitching, into the soluble proteome.

## Introduction

Computational protein design has undergone a revolution in recent years. With the advent of deep learning, the design of proteins with defined structures can now be performed relatively routinely (1, 2). In particular, this is thanks to the development of neural networks for structure and sequence design, and structural prediction (3–6). Together, these methods have enabled the generation of an enormous diversity of protein structures and functions. In short, protein design is no longer solely an academic problem, but has developed into a potentially transformative tool with applications in cell biology, biotechnology, and medicine (7, 8). The newfound success of protein design has opened up an interesting possibility: the design of proteins with functions beyond those found in nature (9). Previously, work focused primarily on the design of *de novo* proteins with non-natural structures (10). However, with the success of deep learning, we can now focus on the more advanced task of designing novel functions.

To expand the functional space available using protein design, we recently turned our attention to the many unique properties found in membrane proteins. We previously reported a deep learning-based method for the design of soluble analogues of membrane proteins (11, 12). With this method, we have demonstrated the design of soluble analogues of several diverse membrane proteins with unique topologies not found naturally in soluble space (12). Moreover, we showed that the soluble analogues can be functionalized with native structural motifs, retaining native protein-protein interactions (PPIs) in the soluble analogues. However, beyond PPIs, there are a huge number of other functions that are only found in membrane proteins, including unique ligand binding modes, switching mechanisms, and dynamic interactions (13, 14). Bringing these into the soluble proteome would add additional functions to the toolbox of molecular and cellular engineering.

One such unique function is retinal-based photoswitching. Retinal is the light-activatable ligand found in the rhodopsin families of photosensors (15). Rhodopsins carry out an enormous range of different functions, ranging from enabling vision in animals, to acting as light-activated enzymes and light-gated ion channels and pumps in bacteria and archaea (16). Rhodopsins have formed the foundation of optogenetics for several decades, enabling the specific control over neuronal activation (17, 18). Previous work has shown that retinal can be incorporated into soluble proteins, and a soluble analogue of bacteriorhodopsin demonstrates that soluble analogues of microbial rhodopsins are tractable (19–21). However, to the best of our knowledge there have been no natural soluble photosensors identified that use retinal as a chromophore. Moreover, there have been no reported soluble analogues of the visual GPCR-fold rhodopsins.

Here, we use the AF2_seq_ framework coupled to the soluble version of ProteinMPNN (ProteinMPNN_sol_), to create soluble analogues of rhodopsins from both the microbial (type I) and animal (type II) families. We modify AF2_seq_ to improve the atomistic design of ligand binding sites in the analogues. We show that the designed soluble analogues retain the characteristic color change of retinal binding to rhodopsin, and confirm the site of retinal binding and Schiff base formation using Top Down mass spectrometry (TD MS). Moreover, X-ray crystallography demonstrates the accurate location of retinal in the designed analogue. Finally, we show that the designed microbial rhodopsin analogue retains the primary reaction of the photocycle, matching very closely the lifetimes of the natural membrane protein under physiological conditions until the early nanoseconds. Together, these data show that unique functions from membrane proteins can be imported into the soluble proteome using AF2_seq_, and suggests a general strategy for designing new functions in soluble proteins.

## Results

### Computational design of soluble rhodopsin analogues

For the computational design of soluble rhodopsin analogues, we used the AF2_seq_-ProteinMPNN_sol_ framework (11). This uses an inverted AlphaFold2 (AF2) network to hallucinate a protein structure, guided towards a target structure by losses (22), followed by a customized version of the ProteinMPNN sequence design network trained only on the soluble proteome (ProteinMPNN_sol_) to create soluble analogues of membrane protein folds (Figure 1A) (4); hereafter referred to collectively as AF2_seq_. AF2_seq_ has been shown previously to be highly proficient at creating soluble analogues of diverse membrane protein folds, even functionalizing the analogues with native motifs to retain natural functions in the soluble analogues. To design soluble analogues of rhodopsins, retinal binding must be retained in the analogues while the transmembrane domains are re-designed. To explore whether AF2_seq_ could be used for this task, we selected three rhodopsins from both the microbial (type I) and animal (type II) families. We chose Sensory Rhodopsin II (*Natronomonas pharaonis*; SRII) from the microbial family, and Bovine Rhodopsin (*Bos taurus*; BovR) and Jumping Spider Rhodopsin-1 (*Hasarius adansoni*; JSR1) from the animal family (23–25). To retain retinal binding, we fixed the residues around the retinal binding pocket, while allowing the rest of the structure to be re-designed (Supplementary Figure 1, Supplementary Table 1). To select the final designs, we re-predicted the designed sequences with AF2 and filtered by confidence of the prediction (predicted value of the local distance difference test (pLDDT) > 80), the surface hydrophobicity (surface apolar fraction (SAF) < 0.2), and the root mean square deviation (RMSD) of the residues coordinating retinal (typically < 1.5 Å).

**Figure 1:**
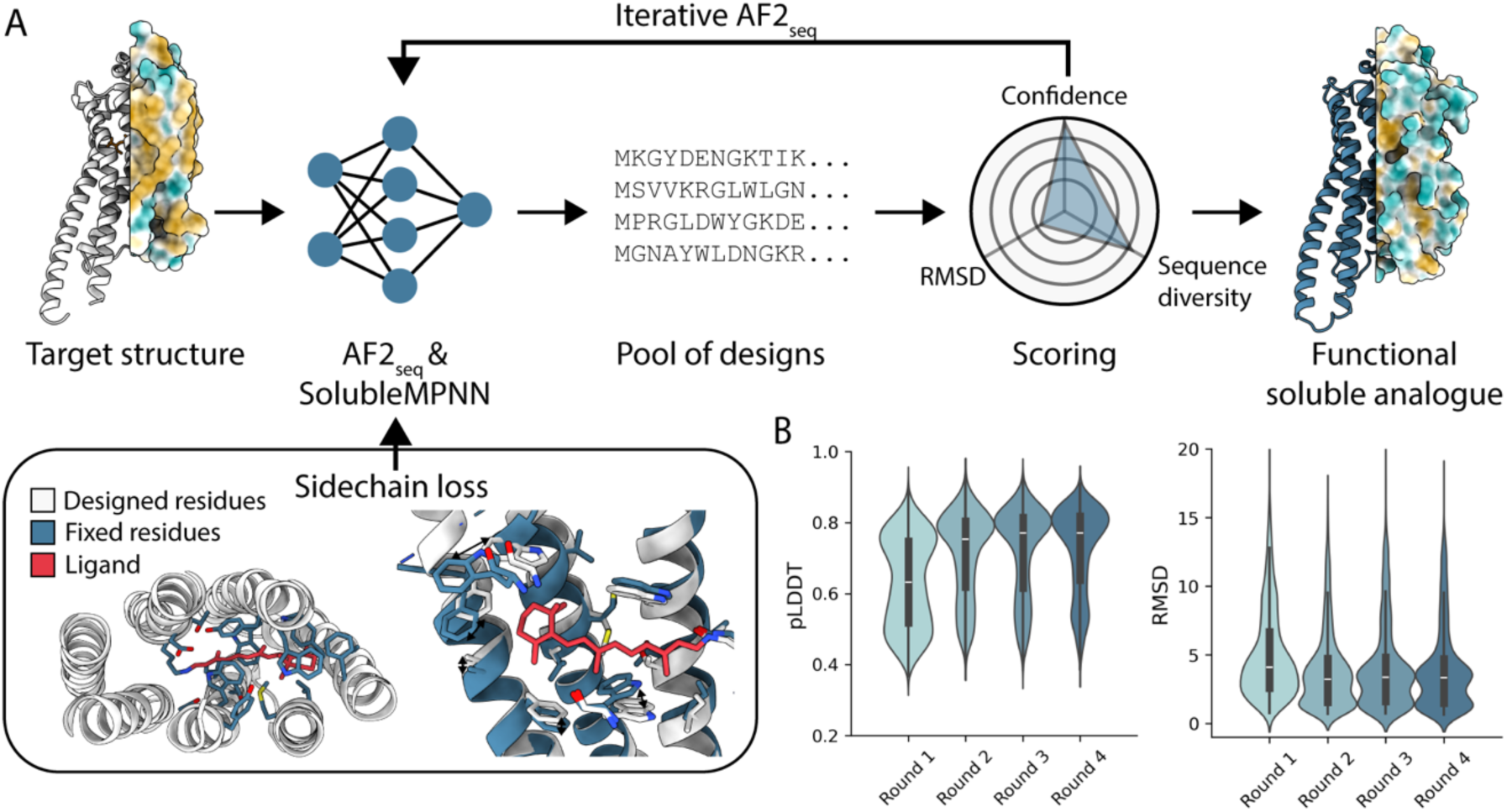
Computational design of soluble rhodopsin analogues. **A** – The AF2_seq_ pipeline for design of soluble analogues of membrane proteins with the additional sidechain RMSD loss for ligand binding. **B** – Reduction in coordinating residue RMSD and increase in pLDDT for the BovR analogue following successive rounds of AF2_seq_ with the applied sidechain RMSD loss.

Using this design method, we observed that there was a large difference *in silico* between the microbial and animal rhodopsins. Running AF2_seq_ on SRII produced high confidence predictions with low RMSDs after a single pass. In contrast, AF2_seq_ struggled to generate sequences passing the *in silico* filters for BovR and JSR1. We hypothesized that this is due to the differing structural complexity of the two targets. The microbial rhodopsins have a relatively simple 7-transmembrane helix fold, while the animal rhodopsins have a G-protein Coupled Receptor (GPCR) fold (16). Moreover, a structural fold similarity search for the three targets on the SCOP database indicated that SRII bears significant structural similarity to soluble proteins (TMscore > 0.5). While, in agreement with other tested GPCR folds, there are no soluble folds similar to BovR or JSR1 (Supplementary Table 2) (12). This reinforced that design of a soluble ligand-binding GPCR analogue may be a more challenging task than for the microbial rhodopsin fold.

To improve our *in silico* success, we implemented two modifications to the structure generation portion of AF2_seq_. Firstly, we implemented an additional sidechain RMSD loss during AF2 backpropagation. This loss minimizes the sidechain RMSD of selected residues during the structure generation trajectory and was applied to residues critical for retinal binding in the animal rhodopsin analogues. Secondly, we found that running several iterations of AF2_seq_, where the new trajectories were initialized from the top scoring sequences from the previous round, generated significantly better scoring designs than after only a single pass (Figure 1B). Using this adjusted pipeline for the animal rhodopsins, we were able to generate sequences for all three targets to experimentally validate.

### Designed soluble analogues of microbial and animal rhodopsins

From AF2_seq_, we ordered sequences for all three targets to evaluate *in vitro*. The soluble rhodopsin analogues were expressed in *Escherichia coli* (*E. coli*) and purified using a *C*-terminal 6-His tag followed by size exclusion chromatography (SEC). Similarly to other soluble analogues generated by AF2_seq_, the majority of the tested proteins were highly soluble and purifiable by size exclusion chromatography (SEC) (SRII: 8 of 12 designs, BovR: 25 of 30, JSR1: 12 of 20).

To evaluate whether the soluble analogues were able to bind retinal, we added an equimolar concentration of retinal to each of the proteins after purification. For SRII we added all-trans-retinal, and for BovR and JSR1 we added 11-cis-retinal, to match the isomer found in the target membrane protein (Figure 2A, B). Retinal binding to rhodopsin is associated with a red-shifting of the absorbance maximum of retinal from 380 nm to typically 440 – 600 nm, depending on the protein environment (26, 27). For our designs, we selected proteins with an absorbance maximum greater than 440 nm (corresponding to the maximum of the protonated retinal Schiff base) as likely binding to retinal (28, 29). Again, SRII showed the greatest success rates, with 4 of 8 expressed designs having an absorbance maximum at or greater than 440 nm (Supplementary Figure 2). BovR also proved to be a tractable design target, with 8 of 25 tested designs showing red shifting at or greater than 440 nm. JSR1 was a more challenging target however, with only 1 of the 12 purified proteins showing any red shifting, with the majority of the retinal still unbound. As the opsin shift is highly dependent on conformational, and electrostatic effects including the solvent environment, the absorption maxima are still blue shifted compared to the membrane protein counterparts (30, 31).

**Figure 2:**
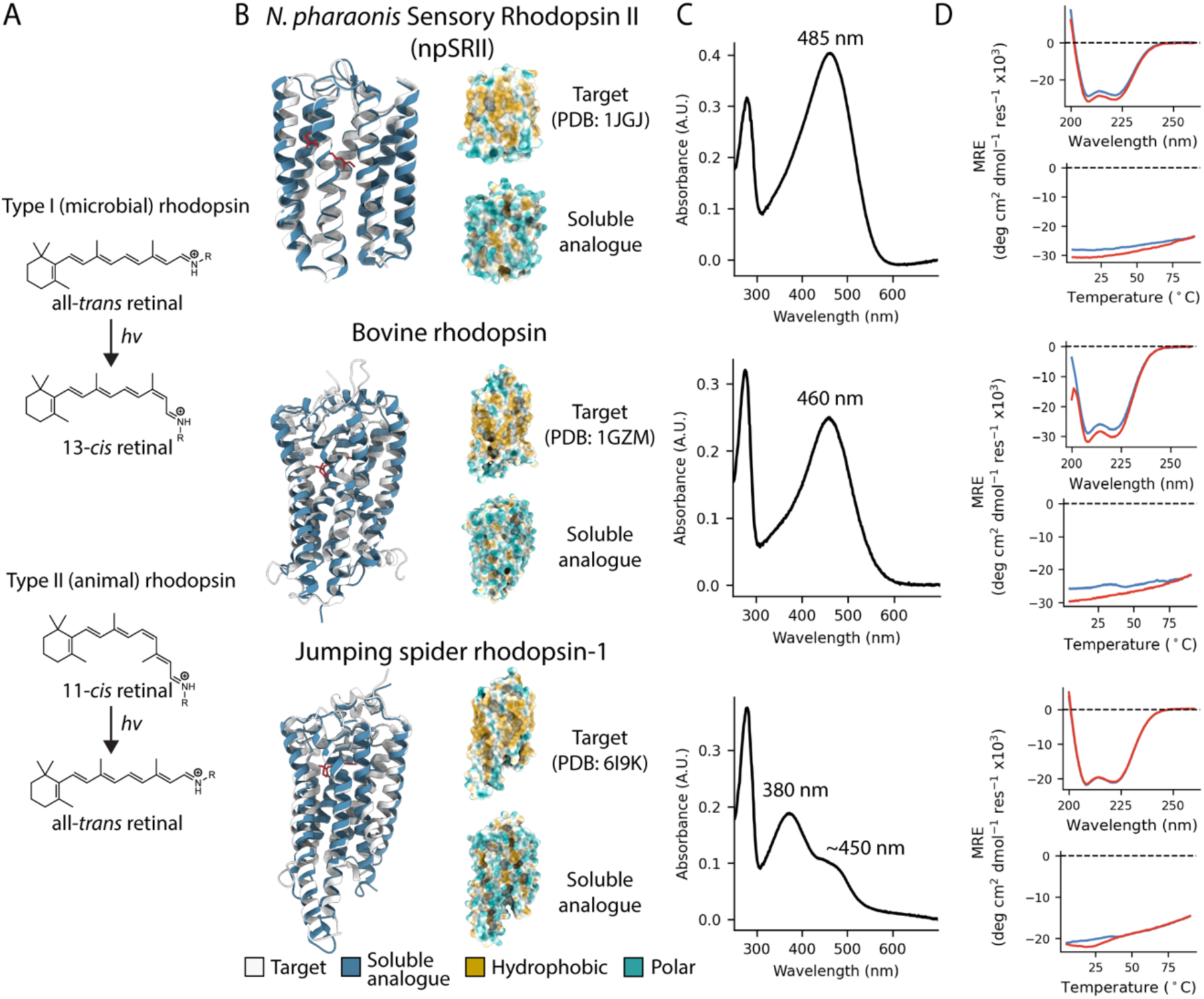
Soluble analogues of microbial and animal rhodopsins. **A** – Predominant retinal isomers found in microbial and animal rhodopsins. **B** – AF2 models of the soluble analogues overlayed with their targets. **C** – Absorbance spectra of the soluble analogues after addition of retinal. **D** – Secondary structure and thermal stability of the soluble analogues as measured by CD spectroscopy.

For each of the three targets, we selected the best design based on the proportion and extent of red shifting for further biophysical characterization (Figure 2C). All of the chosen soluble analogues have below 30% sequence identity to the target (SRII: 24.4%, BovR: 22.4%, JSR1: 28.9%; Supplementary Figure 3). Initially, we evaluated the folding and thermostability of the designs by circular dichroism (CD) spectroscopy. All three soluble analogues showed clear α-helical fingerprints by CD, matching the designed structures (Figure 2D). Moreover, and in agreement with other soluble analogues made by AF2_seq_, the soluble rhodopsins were highly thermostable, with no appreciable denaturation after heating to 90 °C.

### Soluble rhodopsin analogues show precise retinal binding

Having determined that the designs were correctly folded and appeared to show the characteristic spectral signature of retinal binding, we aimed to conclusively determine that retinal was bound in the correct location in the soluble analogue. Firstly, we created point mutations in each of the soluble analogues, mutating the core Schiff base lysine for retinal attachment to alanine. These mutants would not be expected to create a covalent bond to retinal and should not show any red shifting on retinal binding. Indeed, as expected, all three mutants were abrogated for red-shifting, with the soluble SRII analogue (SRII_sol_) and BovR analogue (BovR_sol_) showing absorbance maxima analogous to unbound retinal (380 – 400 nm; Figure 3A). The JSR1 soluble analogue (JSR1_sol_), which previously only showed a small shoulder at approximately 450 nm, now showed a uniform peak with a maximum at 380 nm, confirming that this shoulder had been due to retinal binding.

**Figure 3:**
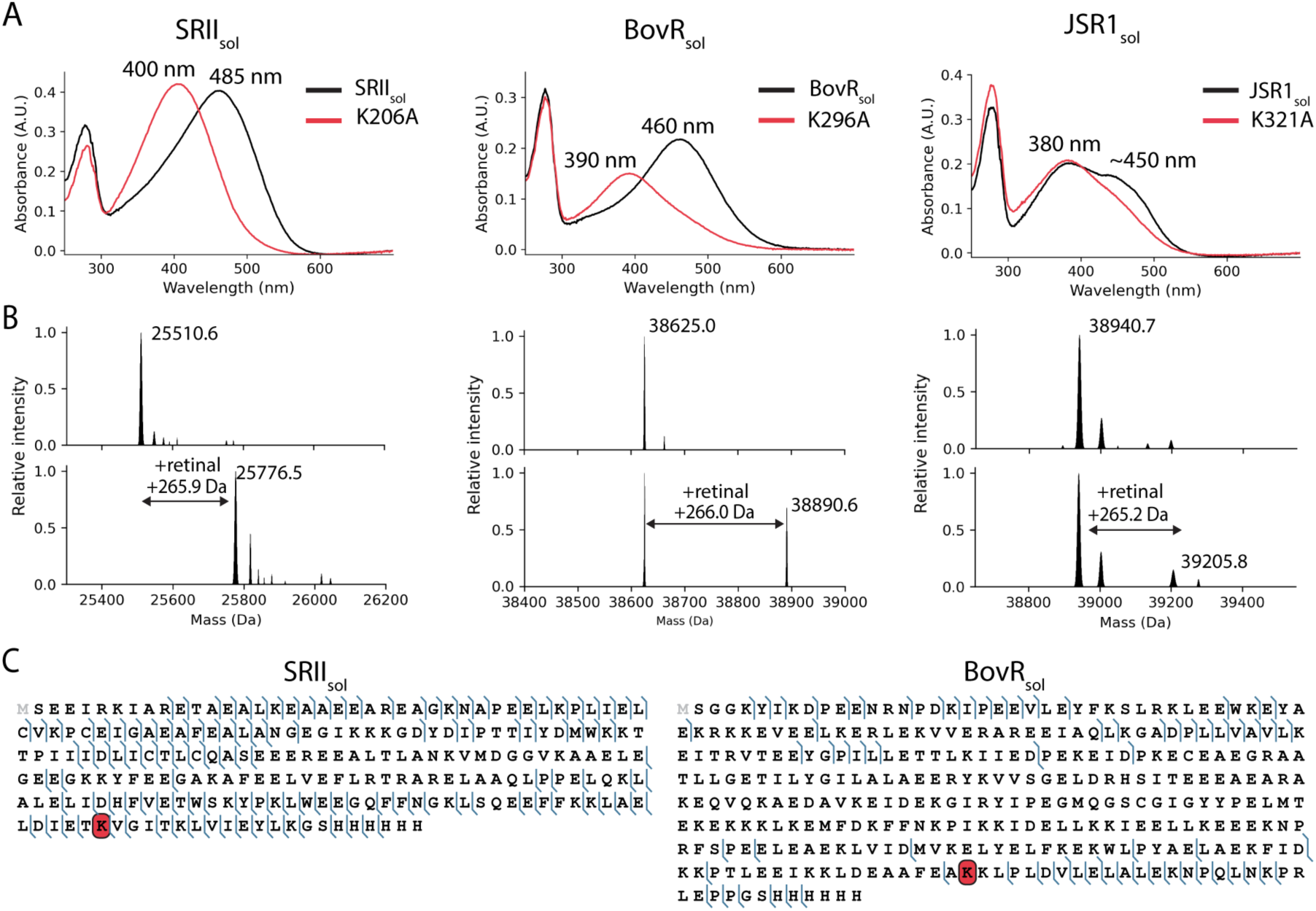
Confirmation of retinal attachment in the soluble analogues. **A** – Absorbance spectra of the soluble analogues and the core lysine mutants. **B** – Native mass spectra of the soluble analogues before and after retinal addition. **C** – Top-down MS fragmentation maps of the soluble analogues with retinal. Detected b- and y-fragment ions are denoted in blue, while retinal modification is marked as a red square on lysine residues.

Secondly, we performed native mass spectrometry (MS) on the protein:ligand complexes. Examination of the soluble analogues by native MS showed in all three a clear mass shift of +266.4 Da after incubation with retinal, corresponding to the expected average mass of retinal in the Schiff base (C_20_H_26_; Figure 3B). Interestingly, and in accordance with the spectroscopy results, SRII_sol_ and BovR_sol_ showed a large amount of protein:ligand complex, indicating efficient formation of the holo protein. However, in JSR1_sol_ only a small fraction of the protein:ligand complex was detected by native MS, as suggested by the relatively poor red shifting.

Finally, to fully confirm the location of retinal attachment, we performed top-down MS on the protein:ligand complexes under denaturing conditions (32, 33). This enables the identification of covalent modifications to the protein residues at a very high mass resolution, allowing detection of residue level modifications depending on the fragmentation pattern. Using this method, each of the soluble analogues was analyzed in turn. The intensity of the retinal-bound proteoform of JSR1_sol_ was insufficient for top-down MS analysis under denaturing conditions. However, both of the SRII_sol_ and BovR_sol_ retinal-bound proteoforms gave assignable fragmentation patterns (Figure 3C). Positioning of the detected fragment ions within the corresponding fragmentation maps confirmed the retinal modification specifically to the fragment containing the designed lysine for Schiff base formation in both SRII_sol_ and BovR_sol_ (SRII_sol_ – K206; BovR_sol_ – K296).

Taken together, these data confirm that the designed soluble rhodopsin analogues bind retinal at the designed location in each case, and with shifts in absorbance maxima similar to the natural membrane proteins.

### Soluble rhodopsin analogues are structurally accurate

To validate the structural accuracy of the soluble rhodopsin analogues in more detail, we solved the crystal structure of SRII_sol_ to 1.76 Å resolution (Supplementary Table 3). Analysis of the structure confirmed its structural fidelity to the target membrane protein: the soluble analogue has a 2.1 Å Cα RMSD to the target, and 2.0 Å Cα RMSD to the AF2_seq_ design model (Figure 4A). The two extracellular anti-parallel β-strands could not be resolved in the structure of the soluble analogue, but there was excellent density for the all-*trans*-retinal in the core of the protein.

**Figure 4:**
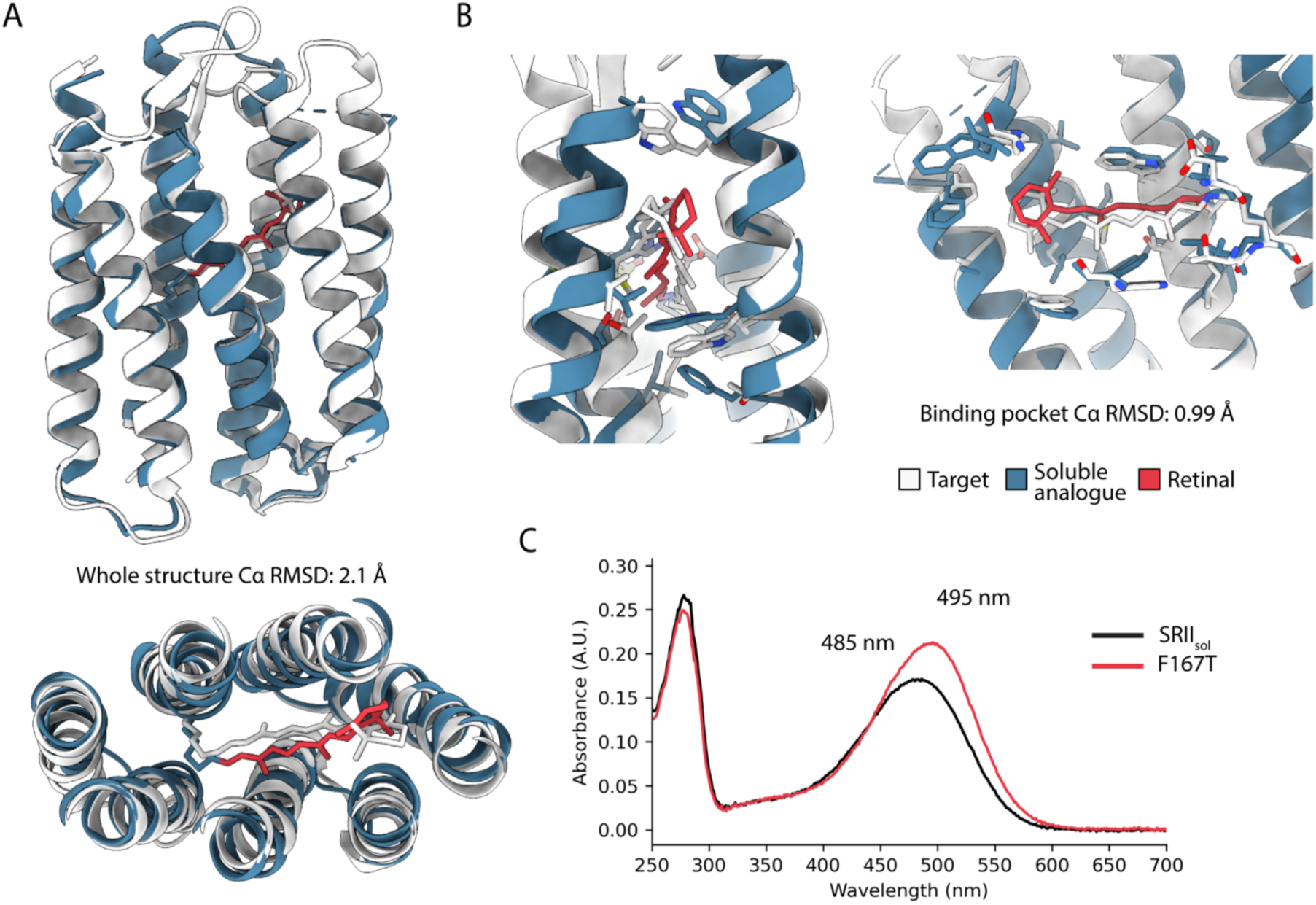
The soluble analogue SRII_sol_ mimics closely the membrane counterpart. **A** – Alignment of the measured structure of the soluble analogue against the target membrane protein. **B** – Alignment of the retinal-coordinating residues in the soluble analogue and the target. **C** – Absorbance spectra of the soluble analogue and the F167T mutant.

Examination of the retinal and the neighboring residues showed again a high level of accuracy in the soluble analogue (0.99 Å Cα/2.3 Å all-atom RMSD to the membrane protein; Figure 4B). Further, the retinal is arranged in a highly consistent way to the target, with the Schiff base clearly formed between K206 and retinal. However, there is a noticeable rotational twist in the retinal in the soluble analogue compared to the target protein, with the chromophore twisted approximately 60° along its axis. Examination of the surrounding residues suggested that this was caused by a slight over-packing of the protein core by AF2_seq_. Here, residue T167 in SRII had not been conserved in the design and had been replaced with a larger Phe residue in the analogue. The resulting larger sidechain appeared to be pushing W171, directly below the center of the chromophore, up into the plane of the retinal methyl groups and forcing a rotational twist. We postulated that fixing this slight overpacking of the core could improve the fidelity of our soluble analogue. Indeed, SRII_sol_-F167T showed a 10 nm red-shifting compared to SRII_sol_, placing its absorption maximum at approximately 495 nm, very close to the maximum reported for SRII (500 nm; Figure 4C) (34).

### SRII_sol_ displays accurate early photocycle intermediates

Finally, we asked whether the designed soluble analogues retained a functional photocycle, analogous to the parent rhodopsins. To this end, we analyzed SRII_sol_-F167T and BovR_sol_ by ultrafast spectroscopy.

We first probed SRII_sol_-F167T. The initial photoreaction of most microbial rhodopsins at physiological pH proceeds from excitation in the ground state via a short-lived electronically excited state and the vibrationally excited J-intermediate to the K-intermediate, which is the last intermediate observed on a 2 ns timescale. The transition to later photointermediates (L/M/N/O) is expected in the micro- to millisecond timespan (35).

Indeed, in SRII_sol_-F167T a strong short-lived excited state absorption (ESA) signal (∼525 nm), which transforms into a more red-shifted signal belonging to the J-intermediate (∼600 nm) and subsequently a slightly less red-shifted signal belonging to the K-intermediate (∼540 nm), is observed (Figure 5A). Their decay and rise lifetimes roughly align with the lifetimes of the natural SRII (36). These data indicate that the primary photoreaction proceeds normally, confirming an intact binding pocket and photoswitching functionality of the retinal. However, at a slower timescale, no other intermediates could be detected, even though the SRII photocycle normally takes up to a second (34). Therefore, the photocycle is either terminated after the shortened K-intermediate, or only minimally populated.

**Figure 5:**
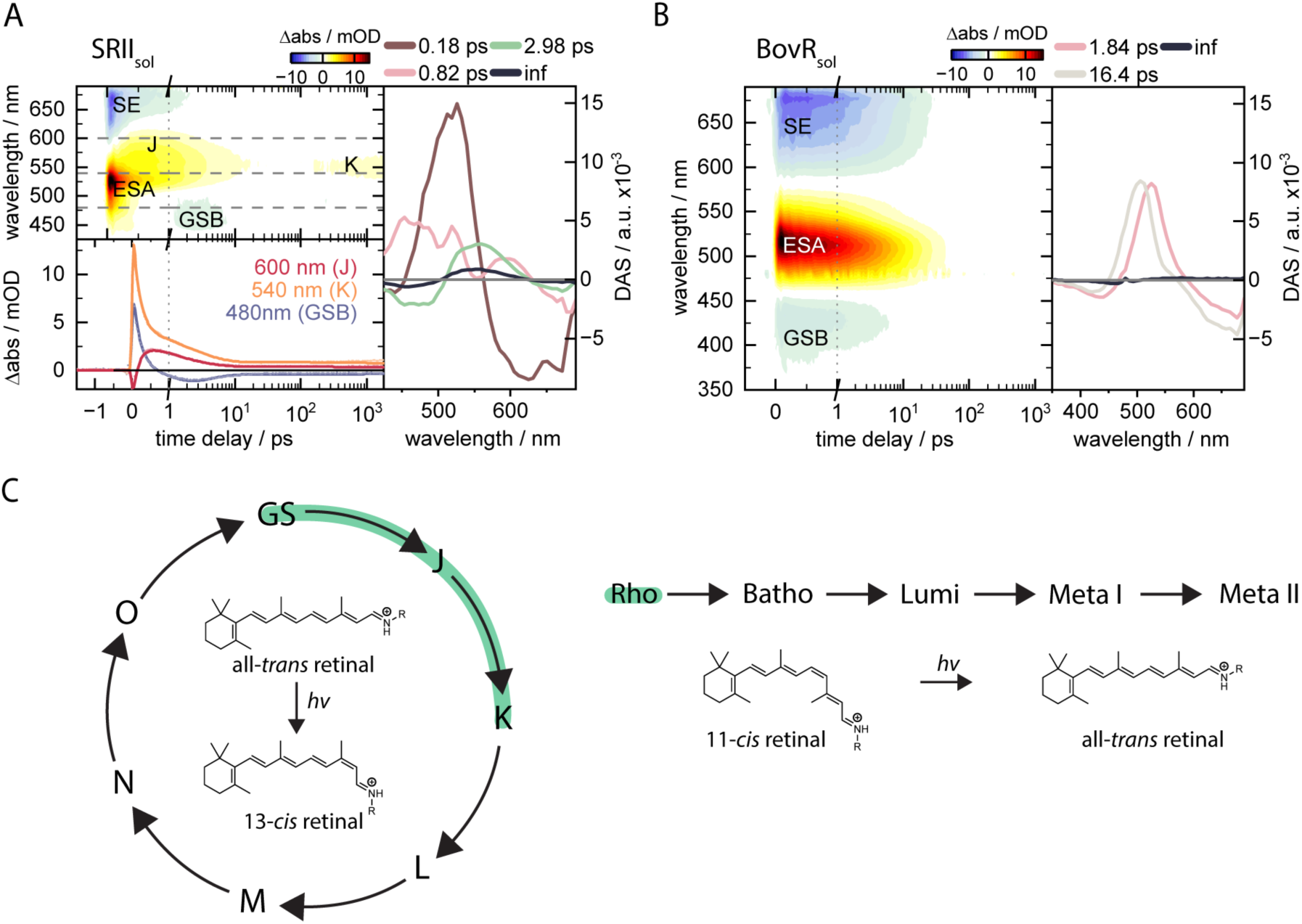
Ultrafast spectroscopy of the soluble rhodopsin analogues. Time-resolved spectroscopy of the initial photoproducts of SRII_sol_ (**A**) and BovR_sol_ (**B**). For each, the excited state absorption (ESA) signal is shown alongside the stimulated emission (SE) and ground state bleach (GSB) signals. For SRII_sol_ the ESA is followed by J- and K-photointermediates. **C** – The predominant photointermediates of natural SRII and BovR, with the observed states in the soluble analogues highlighted in green.

Having validated the early photointermediates in SRII_sol_-F167T, we next examined BovR_sol_. Belonging to the class of animal rhodopsins, BovR is expected to display several intermediates in the femto- to nanosecond range: photorhodopsin at ∼570 nm, bathorhodopsin at ∼540 nm and a blue shifted intermediate at ∼480 nm (37). However, instead of the anticipated fast formation of the primary photoproducts of the BovR photocycle, a long-lived ESA and stimulated emission (SE) represent a hindered retinal isomerization. Both signals and the ground state bleach (GSB) decay with two lifetimes of 2 and 16 ps, leaving no residual signal (Figure 5B). Overall, these represent a restricted chromophore in BovR_sol_, with no visible photointermediates.

## Discussion

In summary, we have designed soluble analogues of both microbial and animal rhodopsins using AF2_seq_. This task involved developing AF2_seq_ for the design of ligand binding pockets, a task for which it had not previously been applied. For this, we developed an additional sidechain RMSD loss and used successive iterations of AF2_seq_. The tested designs show a high fidelity to the target membrane proteins, with accurate chromophore binding and high structural similarity. Moreover, we have demonstrated that the soluble analogues can mimic the natural functions of rhodopsins, with the primary photoreaction of SRII_sol_ matching the natural protein under physiologically relevant conditions. We note that only the initial steps of the photocycle are retained in SRII_sol_ (Figure 5C) (36). As microbial rhodopsin photocycles can be disrupted by mutation of only a few key amino acids, some key residues for the later photointermediates are most likely missing in the analogue. This likely results in destabilization of the K-intermediate resulting in a shorter lifetime, preventing conformational rearrangements of the protein to other intermediates. This could also be due to the overall rigidity of the binding pocket in the soluble analogues. This is even more evident in BovR_sol_. Here the rigidity of the retinal binding pocket likely impedes the formation of even early photointermediates. This could be in part due to the larger structural rearrangements found in natural BovR, which have not been included in the design trajectory and would require designing for multiple retinal conformations.

Previous work showed that ProteinMPNN_sol_ alone was able to create soluble analogues of the microbial bacteriorhodopsin (21). Here, we have shown that AF2_seq_ can not only produce analogues of microbial rhodopsins with a high fidelity, but also produce analogues of the more complex animal rhodopsins. These targets are more challenging not only because the GPCR fold does not exist in the soluble proteome, but also because the animal rhodopsins have a much greater level of structural dynamics compared to the microbial rhodopsins (38). Moreover, all of the designed soluble analogues have a significantly lower sequence identity to the membrane protein than has been shown previously, and are accompanied by a high thermostability. We note also that the designed SRII_sol_ has an absorbance maximum very close to the design target, and retains early photointermediates that match the native photocycle at physiological pH, which was not possible using ProteinMPNN_sol_ alone (21).

Overall, this reinforces the generalizability of the AF2_seq_ framework, and its capability to design analogues that are not only structurally faithful to the target, but can incorporate function as well. We envisage that in future this could be applied to diverse ligands, particularly considering the range of natural ligand-binding pockets found in GPCRs. Soluble ligand-binding GPCRs could unlock a range of small molecule sensors with high stability and facile expression, ideal for synthetic biology applications.

More broadly, this work advances the possibility to bring unique membrane protein functions into the soluble proteome. To the best of our knowledge, there have been no natural proteins identified to use retinal as a chromophore in the soluble proteome. This could open the door to the design of novel soluble photoresponsive proteins that use retinal as a chromophore. Other unique membrane protein functions could also be explored for solubilization, including unique methods of switching found in transport proteins and channels. Despite the progress on the design of membrane soluble analogues, we gather that we are still missing the incorporation of conformational dynamics found in membrane proteins, which remains unclear if it is achievable by computational design

## Materials and Methods

### Structural fold similarity search

The fold similarity search was performed using FoldSeek on the SCOP database (downloaded March 2023) separated into globular and membrane proteins using the domain annotations (39). For each of the target folds, an exhaustive search on the basis of TM score alignment was performed where a match was defined as TM > 0.5 (40).

### Computational design

A full end-to-end pipeline for the computational design of soluble analogues of membrane proteins is provided at https://github.com/alexhilditch/solubilization. This script follows the AF2_seq_-ProteinMPNN_sol_ design method within the ColabDesign framework, enabling all of the steps (structure-sequence hallucination, sequence design, and structural prediction) to be performed with minimal intervention.

Here, AF2_seq_ was used to design soluble analogues of the three target rhodopsin structures of SRII (PDB: 1JGJ), BovR (PDB: 1GZM), and JSR1 (PDB: 6I9K). The *N*-terminal 19 residues and *C*-terminal 30 residues of JSR1 were omitted from the design trajectory as they are not resolved in the target structure, however residue numbering throughout the text is consistent with JSR1 for clarity.

Computational design was performed in three stages: 1) coupled backbone-sequence generation using AF2_seq_, 2) sequence design with ProteinMPNN_sol_, and 3) re-prediction and filtering with AF2. In each case, residues important for retinal binding were fixed throughout the design pipeline. Each AF2_seq_ design trajectory used 500 rounds of gradient descent optimization, creating 50 backbones per design. The AF2_seq_ network was constructed as described previously, with the addition of a sidechain loss for optimization of ligand binding. All design runs were executed on a single Nvidia Tesla H100 (94 GB) GPU.

Previously, AF2_seq_ used a compound loss function for the computation of error gradients:

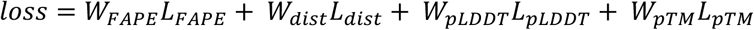

Where *W* and *L* are the weight and value of the loss respectively. The frame aligned point error (FAPE) loss quantifies the L2 norm between the predicted Cα atoms and the target structure. The distogram (*dist*) loss is the cross entropy over the Cβ distogram (Cα for glycine). The model confidence (*pLDDT*) loss is given by 1 − pLDDT, penalizing low confidence. The pTM loss is an additional prediction confidence metric focused on global structural similarity.

For the design of SRII, these losses were sufficient to create designs that passed the described in silico filters using AF2 in single sequence mode (without multiple sequence alignments or templates). However, for BovR and JSR1, there were few to no convergent trajectories. To improve this, an additional sidechain loss was implemented. This loss was constructed as an all-atom distogram loss (*distAA*), given by the cross entropy over all atom positions for selected residues, rather than only the Cβ. This loss was applied selectively to the positions in the ligand binding pocket to improve the resolution of the design trajectory in these positions, creating a sidechain-aware design trajectory in specified positions.

In addition, to resolve the non-convergence of the trajectories, trajectories for BovR and JSR1 were ‘soft-started’ from top-ranking sequences from the previous trajectories. Here, the top 15 – 20 sequences from a design round were used as the starting point for further trajectories, in the case of BovR up to four times, improving the number of passing designs from 0 to 25.

From AF2_seq_, 50 sequences were generated with ProteinMPNN_sol_ per backbone, and the resulting sequences predicted with AF2. Where possible, AF2 was used in single sequence mode, but for JSR1 partial templates were required as the intracellular and extracellular parts of the GPCR fold could not be predicted in single sequence mode.

### Protein expression and purification

DNA fragments (Twist Bioscience) encoding for the designed soluble analogues were cloned into pET-11 derivative vectors using either Gibson assembly or golden gate assembly to produce sequences with a *C*-terminal 6xHis tag. The sequence verified vectors were transformed into BL21 DE3 *E. coli* and grown (37 °C, 220 rpm) overnight in lysogeny broth (LB). 250 ml LB expression cultures were inoculated 1:100 with overnight culture and grown (37 °C, 220 rpm) to optical density (OD) 0.4 – 0.6, before inducing with 500 mM Isopropyl β-D-1-thiogalactopyranoside (IPTG) and growing overnight (18 °C, 220 rpm). The cells were collected by centrifugation (5000 xg, 15 minutes) and resuspended in 30 ml lysis buffer (500 mM KCl, 50 mM HEPES pH 7.5, 15 mM imidazole, 1 mg/ml DNase, 1 mg/ml lysozyme, 1mM PMSF). The resuspended pellets were lysed by sonication and clarified by centrifugation (20,000 xg, 20 minutes). The clarified lysate was applied to a 0.5 ml Ni-NTA Superflow column (Qiagen) and washed with 10 column volumes of wash buffer (500 mM KCl, 50 mM HEPES pH 7.5, 15 mM imidazole), before elution in 2 column volumes of elution buffer (250 mM KCl, 50 mM HEPES pH 7.5, 500 mM imidazole). The elute was then applied directly onto a Superdex S200 10/600 or Superdex S75 10/600 gel filtration column (Cytiva) in 150 mM KCl, 50 mM HEPES pH 7.5. The eluted protein was concentrated, frozen in liquid nitrogen, and stored at -80 °C.

### Measurement of retinal binding

Retinal stocks at 1 mM were prepared in the dark by dissolving 11-cis or all-trans retinal in ethanol and concentration determined by measurement of absorbance at 380 nm using the literature extinction coefficients (11-*cis*: 24,935 M^-1^cm^-1^, all-*trans*: 42,880 M^-1^cm^-1^) (41). Stocks were stored at -80 °C. For determination of retinal binding to the soluble analogues, 40 mM of the soluble analogue in 150 mM KCl, 50 mM HEPES pH 7.5 was incubated with 30 mM of the corresponding retinal isomer (SRII: all-*trans*, BovR & JSR1: 11-*cis*) in a black clear-bottomed 96-well plate, in the dark at 4 °C overnight. An absorbance spectrum was then measured using a Tecan Safire 2 plate reader measuring from 240 nm to 700 nm in 1 nm increments. Hits were selected as designs that had an absorbance maximum or visible shoulder at 440 nm or greater.

#### Circular dichroism (CD) spectroscopy

CD spectra were measured using a Chirascan V100 CD spectrometer with a 1 mm path length cuvette. Proteins were measured at 5 mM in 100 mM KCl, 10 mM HEPES pH 7.5. Full spectra measured ellipticity between 200 nm and 260 nm in 1 nm intervals, with a scanning speed of 20 nm min−1 and a response time of 0.125 s. A reference spectrum using the same parameters and buffer at 5 °C was subtracted from the measured ellipticity. Measurements of ellipticity with respect to temperature (melting and cooling spectra), were recorded by measuring ellipticity at 222 nm every 1 °C from 5 – 90 °C and the reverse. Pre-melt and post-melt full spectra were collected before heating, and after cooling respectively. Ellipticity (deg) values were converted to mean residue ellipticity (MRE) (deg·cm2·dmol-1·res-1) by normalization to the number of peptide bonds in the protein, and the path length using the following equation:

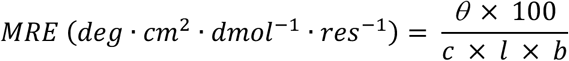

Where *θ* is the difference in absorbed circularly polarized light in millidegrees, *c* is the protein concentration in mM, *l* is the path length in cm, and *b* is the number of amide bonds in the protein.

### Mass spectrometry

#### Intact mass analysis in native-like conditions

Samples were diluted with 50 mM ammonium acetate or injected as they were in MilliQ water with injection volume of 2 µl. The separation was performed on LC instrument (Vanquish Horizon, Thermo Fisher Scientific) using the NativePac OBE-1 SEC column (2.1 x 50 mm, 3 µm, Thermo Fisher Scientific) with isocratic flow of 50 mM ammonium acetate with flow rate 100 µl/min and column oven heated at 30 C. Eluting proteoforms were analyzed using Exploris 240 Orbitrap FT-MS instrument operated in the intact protein mode with High Pressure option activated. Mass spectra were acquired as full scans in the range 1500-8000 m/z at a resolution set to 15 000, RF Lens at 200 %, normalized AGC target of 300 %, maximum injection time of 200 ms, SID of 50 eV, while 10 microscans were averaged. The mass spectra were deconvolved using Biopharma Finder 5.3 (Thermo Scientific) with Respect algorithm

#### Intact mass analysis in denaturing conditions

Samples were injected as they were in MilliQ water with injection volume of 2 µl. The separation was performed on LC instrument (Vanquish Horizon, Thermo Fisher Scientific) using Aquity UPLC Protein BEH C4 pre-column (2.1 x 5 mm, 1.7 µm, 300 A, Waters) with flow rate of 400 µl/min and oven temperature set to 45 °C. The following gradient of solvent B was used: from 15 to 50 % within 2.5 min followed by column washing and re-equilibration steps. Solvent A was composed of MilliQ water with 0.1% formic acid, while solvent B consisted of acetonitrile with 0.1% formic acid. Eluting proteoforms were analyzed in positive polarity using Exploris 240 Orbitrap FT-MS instrument operated in the intact protein mode with Low Pressure option activated. Mass spectra were acquired as full scans in the range 700-4000 m/z at a resolution set to 15 000, RF Lens at 150 %, normalized AGC target of 300 %, maximum injection time of 200 ms, SID of 15 eV, while 10 microscans were averaged. The mass spectra were deconvolved using Biopharma Finder 5.3 (Thermo Scientific) with Respect algorithm.

#### Top-down MS analysis in denaturing conditions

Here, the soluble analogues were analyzed by liquid chromatography (LC) high-resolution MS/MS at a number of different higher-energy collisional dissociation (HCD) energies to fragment the protein and produce fragment ions. The resulting high resolution MS/MS spectra were deconvolved and the obtained neutral masses of b- and y-fragment ions were pooled together. Experimentally detected fragments were matched against the theoretical fragments of the designs including the addition of theoretical ones resulting in +266.2 Da (monoisotopic mass of retinal in the Schiff base, C20H26).

To localize the modification sites, each sample was submitted to top-down MS analysis using higher energy collision induced dissociation (HCD). LC conditions were exactly the same as the ones used for intact mass analysis in denaturing conditions described above. Eluting proteoforms were analyzed on an Orbitrap Exploris 240 FT-MS benchtop instrument (Thermo Fisher Scientific, Bremen, Germany) operated in the intact protein mode with Low Pressure option activated, in positive polarity and using targeted MS/MS approach with ion multiplexing. For this, six precursor ions corresponding to different charge states of the same proteoform carrying retinal modification were input into the targeted inclusion mass list. The isolation window was set to 1.2 m/z, AGC target set as standard, maximum IT set as Auto, with 240’000 resolution at 200 m/z and averaging 10 microscans. Top-down MS analysis was repeated using 6 different values (between 20-50 %) of normalized collision energy (NCE). Top-down MS data were processed using Peak-by-Peak software with MS deconvolution workflow (Spectroswiss, Lausanne, Switzerland).

#### X-ray crystallography

To prepare SRII_sol_ for X-ray crystallography, and due to the limited solubility of retinal in aqueous buffers, the apo-protein was first incubated at 40 mM with 40 mM all-trans retinal in 150 mM KCl, 50 mM HEPES pH 7.5 overnight. Unbound retinal was then removed by desalting before concentration of the holo-protein to 18 mg/ml. Crystals were grown in the dark at 18 °C using the sitting-drop vapor-diffusion method in drops containing 200 nl of protein complex and 200 nl of reservoir solution containing 0.2 M sodium acetate pH 4.5, 30% PEG smear low. For cryoprotection, crystals were briefly immersed in mother liquor containing 20% ethylene glycol. Diffraction data were recorded with ID30A-1 at the European Synchrotron Radiation Facility. The diffraction data were integrated and processed to 1.76 Å by AutoProc. The crystals belonged to space group P 1 21 1. The structure was determined by molecular replacement using PHENIX Phaser, with the AF2 model of SRII_sol_ as a search model. Manual model building was performed using Coot and automated refinement was performed using PHENIX Refine.

#### Ultrafast transient absorption (TA) spectroscopy

Ultrafast TA spectroscopy experiments were performed using a home-built pump-probe setup described in detail elsewhere (42). A Ti:Sa chirped pulse regenerative amplifier (MXR-CPA-iSeries, Clark-MXR Inc., USA) served as the fs-laser source for the setup. It was operated at a central wavelength of 775 nm and a repetition rate of 1 kHz, resulting in laser pulses with a pulse width of 150 fs. Excitation pulses were generated from a home-built two-stage noncollinear optical parametric amplifier (NOPA) with an interlayed prism compressor. For probing absorbance changes, the laser fundamental was focused into a 5 mm CaF2 crystal to generate supercontinuum pulses. The probe light was then divided and guided through sample and reference paths. Two identical spectrographs (Multimode, AMKO, Germany), equipped with a grating (500 nm blaze, 1200 grooves per mm), a photodiode array (S8865-64, Hamamatsu Photonics, Japan) and a driver circuit (C9918, Hamamatsu Photonics, Japan) were used for signal detection. The obtained signals were digitized via a 16 bit data acquisition card (NI-PCI-6110, National Instruments, USA). The pump wavelength was adjusted to the respective absorption maximum, and pump and probe pulses were set to the magic angle (54.7°) configuration to eliminate anisotropic effects. Additionally, the sample was constantly moved in a plane perpendicular to the excitation beam to avoid multifold excitation and sample degradation. Samples were prepared in a 50 mM HEPES, 150 mM KCl buffer at pH 7.5 in a 1 mm cuvette. Absorption spectra were taken before and after each experiment to verify sample quality.

## Acknowledgments

We thank the National Eye Institute (NEI) for providing 11-*cis*-retinal. We thank the developers of the ColabDesign protein design developers (Sergey Ovchinnikov, Shihao Feng, Justas Dauparas, Weikun Wu, and Christopher Frank). We thank F. Pojer, K. Lau and A. Larabi (Protein Production and Structure Characterization Core Facility, EPFL, Switzerland) for help with crystal screens and data collection. A.T.H. was supported by the Peter und Traudl Engelhorn Stiftung Postdoctoral Fellowship. We thank the Swiss National Science Foundation (SNSF) for funding and support.

## Data and code availability

The crystal structure of SRII_sol_ has been deposited in the protein data bank (PDB) under accession code 29GM. Design metrics and models of the tested soluble analogues are accessible on Zenodo (https://doi.org/10.5281/zenodo.20926345). Code for the original implementation of AF2seq can be found at https://github.com/bene837/af2seq. Code for the ColabDesign framework can be found at https://github.com/sokrypton/ColabDesign. An end-to-end pipeline for the design of a soluble analogue of a membrane protein can be found at https://github.com/alexhilditch/solubilization.

## Author Contributions

All authors were involved in conceiving the project and designing the experiments. A.T.H. and C.A.G. created the computational pipeline and created the designs. A.T.H., N.G., S.W., J.W., and L.M. designed and performed the wet lab experiments. All authors contributed to data analysis and preparation of the manuscript.

