## Supplementary Information for "Computational design of microbial and animal rhodopsin soluble analogues"

### Supplementary Figures and Tables

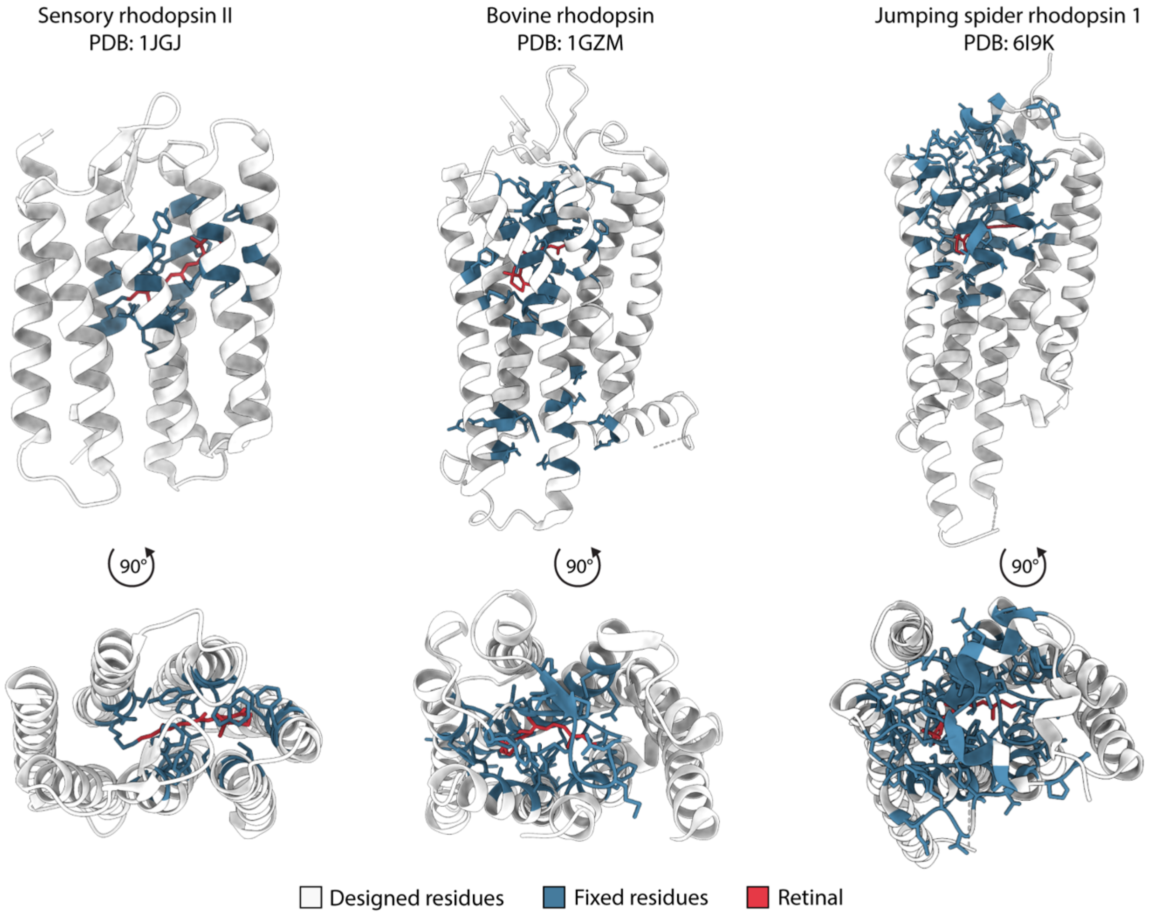


**Supplementary Figure 1: Designed and fixed positions.** The residues fixed during design with $AF2_{seq}$ are colored in blue, residues redesigned are colored in white.

| Target | Fixed positions |
|---|---|
| Sensory rhodopsin II | 73,76,79,80,83,108,109,112,127,130,131,134, 171, 174, 175,178,201,204,205 |
| Bovine rhodopsin | 72,76,110,113,114,117,118,121,122,125,134, 135, 136,138,139,178,179,180,181,182,183, 184,186, 187, 188,189,191,192, 207,208,212, 246,247,250, 253, 261,265,268, 269,272,292,293,295,296,310,312 |
| Jumping spider rhodopsin 1 | 31,33,34,35,36,37,38,39,41,43,44,103,116, 119,120, 123,124,126,127,130,131,134,135, 138,190,191,192, 193,194,195,196,197,198, 199,200,201,202,203, 204, 205,206,207,211,214,215,218,219,222, 223,226,286, 289,290,291,292,293,294,297, 298,300,301,303,304, 305,306,308,309,310, 313,314,316,317,318,320,321 |

**Supplementary Table 1: Fixed positions during AF2seq design.** Residues are numbered according to the parent membrane protein for consistency.

| Query PDB | Target PDB | TM score |
|---|---|---|
| 1JGJ (SRII) | 2P7V_A_1_158 | 0.5881 |
| | 2GSC_A_9_125 | 0.559 |
| | 2JX0_A_640_770 | 0.5334 |
| | 2JX0_A_640_770 | 0.5305 |
| | 2RLD_C_1_115 | 0.5229 |
| 1GZM (BovR) | 2LEM_A_1_216 | 0.4461 |
| | 2LEM_A_1_216 | 0.445 |
| | 1DKZ_A_507_603 | 0.4445 |
| | 2B0H_A_1843_1973 | 0.4442 |
| | 2LEM_A_1_216 | 0.4437 |
| 6I9K (JSR1) | 2LEM_A_1_216 | 0.4604 |
| | 2B0H_A_1843_1973 | 0.4584 |
| | 2LEM_A_1_216 | 0.4576 |
| | 2LEM_A_1_216 | 0.4555 |
| | 2B0H_A_1843_1973 | 0.4548 |

**Supplementary Table 2: FoldSeek search of structural similarity.** The top 5 results by TMscore for each membrane protein target against proteins in the SCOP database labelled as soluble proteins.

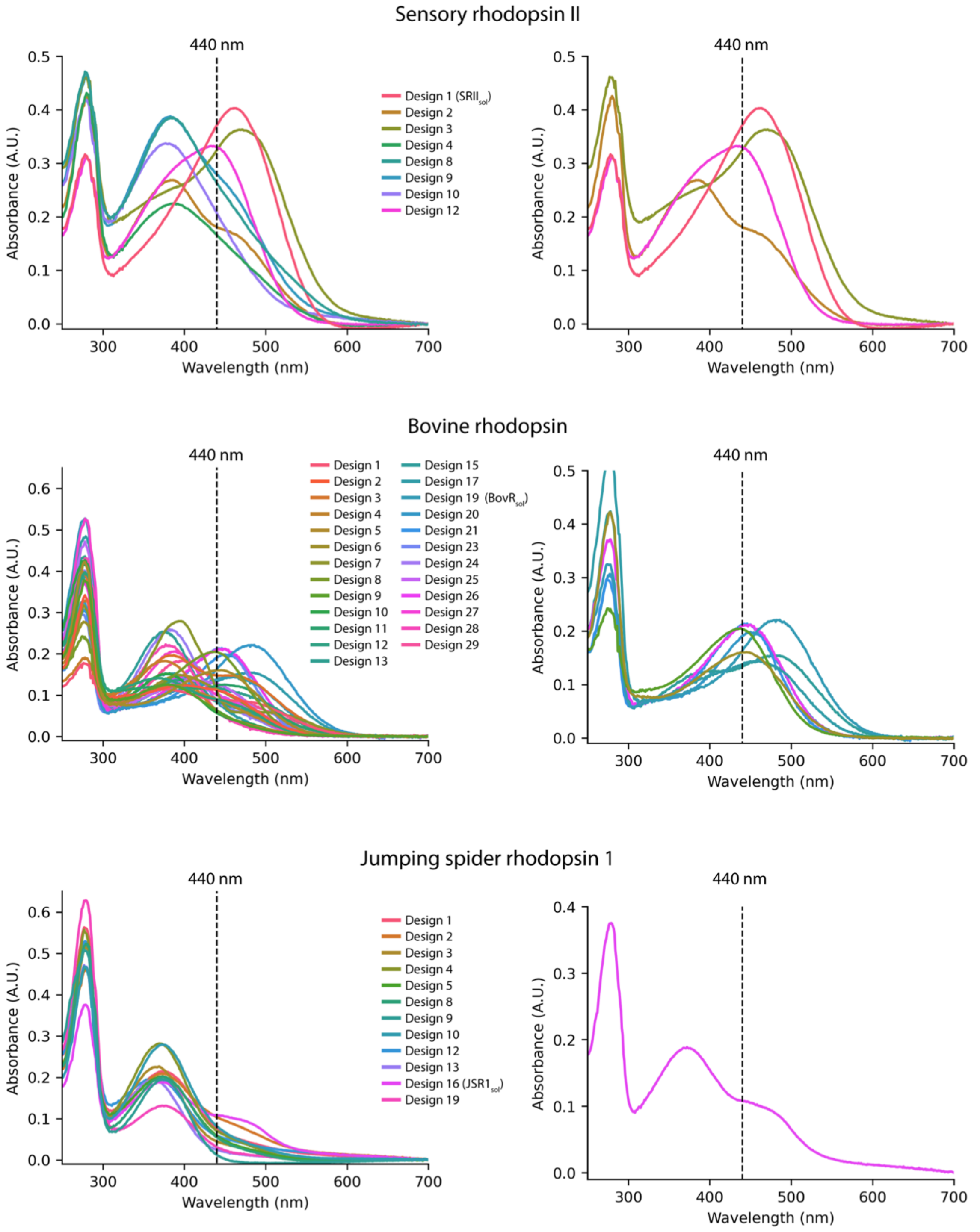


**Supplementary Figure 2: Screening for retinal binding.** Absorbance scan for each of the purifiable designs after incubation with retinal. **Left**: All measured designs. **Right**: Designs with an absorbance maximum or shoulder at or greater than 440 nm.

```
SRII_sol    1 S E E I R K I A R E T A E A L K E A A E E A R E A G K N A P E E L K P L I E L C V K P C E I G A E A
              . . . : . . : . . . . | . . : . . . . . . . . . | | : : | . . . . : . . . . . . | . . . . | . | . |
SRII        1 M V G L T T L F W L G A I G M L V G T L A F A W A G R D A G S G E R R Y Y V T L V G I S G I A A V A

SRII_sol   51 F E A L A N G E G I K K K G D Y D I P T T I Y D M W K K T T P I I I D L I C T L C Q A S E E E R E E
              : . . : | . | . | . . . . . : . . : . . . . | . . | . . | | | : | : . . : . . | . . . . . . | . . .
SRII       51 Y A V M A L G V G W V P V A E R T V F V P R Y I D W I L T T P L I V Y F L G L L A G L D S R E F G I

SRII_sol  101 A L T L A N K V M D G G V K A A E L E G E E G K K Y F E E G A K A F E E L V E F L R T R A R E L A A
              . : | | . . . | | . . | . . . | . : . | . | . . . . | . . | | . | | . . | | . : | . . . . . | . | :
SRII      101 V I T L N T V V M L A G F A G A M V P G I E R Y A L F G M G A V A F I G L V Y Y L V G P M T E S A S

SRII_sol  151 Q L P P E L Q K L A L E L I D H F V E T W S K Y P K L W E E G Q F F N G K L S Q E E F F K K L A E L
              | . . . . : : . | . : . | . : . . | . . | : . | | . : | . . | . . . . . . . | : . . . . . . . : . . |
SRII      151 Q R S S G I K S L Y V R L R N L T V V L W A I Y P F I W L L G P P G V A L L T P T V D V A L I V Y L

SRII_sol  201 D I E T K V G I T K L V I E Y L K
              | : . | | | | . . . : . : : . . .
SRII      201 D L V T K V G F G F I A L D A A A
```

**Identity:** 53/217 (24.4%)
**Similarity:** 82/217 (37.8%)
**Gaps:** 0/217 (0.0%)

```
BovR_sol    1 G K Y I K D P E E N R N P D K I P E E V L E Y F K S L R K L E E W K E Y A E K R K K E V E E L K E R
              . . . . : . | . . . . . . . . . . . . | . . . | : : . : . . . . . . . . . . . . . . . : . . | . . .
BovR        1 M N G T E G P N F Y V P F S N K T G V V R S P F E A P Q Y Y L A E P W Q F S M L A A Y M F L L I M L

BovR_sol   51 L E K V V E R A R E E I A Q L K G A D P L L V A V L K E I T R V T E E Y G P I L L E T T L K I I E D
              . . . : . . . . . . . . . | . | . . . . . | . . : | . . : . . . . . . . . . . . . . | | | . . . . .
BovR       51 G F P I N F L T L Y V T V Q H K K L R T P L N Y I L L N L A V A D L F M V F G G F T T T L Y T S L H

BovR_sol  101 P E K E I D P K E C E A E G R A A T L L G E T I L Y G I L A L A E E R Y K V V S G E L D R H S I T E
              . . . . . . | . . | . . | | . . | | | . | | . . | : . : : . | | . | | | . | | . . . . . : . . . . |
BovR      101 G Y F V F G P T G C N L E G F F A T L G G E I A L W S L V V L A I E R Y V V V C K P M S N F R F G E

BovR_sol  151 E E A E A R A K E Q V Q K A E D A V K E I D E K G I R Y I P E G M Q G S C G I G Y Y P E L M T E K E
              . . | . . . . . . . . . . | . . . . . . . . . . . . | | | | | | | | . | | | | . | | . . . . . . . .
BovR      151 N H A I M G V A F T W V M A L A C A A P P L V G W S R Y I P E G M Q C S C G I D Y Y T P H E E T N N

BovR_sol  201 K K K L K E M F D K F F N K P I K K I D E L L K K I E E L L K E E E K N P R F S P E E L E A E K L V
              : . . : . . | | . . . | . . | : . . | . . . . . : : . . . : | | . . . . . : . | . . . . : | | | . |
BovR      201 E S F V I Y M F V V H F I I P L I V I F F C Y G Q L V F T V K E A A A Q Q Q E S A T T Q K A E K E V

BovR_sol  251 I D M V K E L Y E L F K E K W L P Y A E L A E K F I D K K P T L E E I K K L D E A A F E A K K L P L
              . . | | . . : . . . | . . . | | | | | . : | . . . . . . : . : . . . . . . : . . . | | . | | . . . :
BovR      251 T R M V I I M V I A F L I C W L P Y A G V A F Y I F T H Q G S D F G P I F M T I P A F F A K T S A V

BovR_sol  301 D V L E L A L E K N P Q L N K P R L E P P
              . . . . : . : . . | . | . . . . . : . . .
BovR      301 Y N P V I Y I M M N K Q F R N C M V T T L
```

**Identity:** 72/321 (22.4%)
**Similarity:** 101/321 (31.5%)
**Gaps:** 0/321 (0.0%)

```
JSR1_sol    1 K M I D T V T D D M K P M I H E H W K K F P P L P E E V Y E E I K K I G E E I E K K I E E I K K K L
              . : : | . : . : | | . | | | | | | | . | | | | : . . . . : : . . : . . : . . . | . . . . . . . . . :
JSR1        1 S I V D L L P E D M L P M I H E H W Y K F P P M E T S M H Y I L G M L I I V I G I I S V S G N G V V

JSR1_sol   51 E E I L K S K E L D P N D P L S F Q A K K L L E F I E K I E E M L K L E N G E E G K N T W H K G P K
              . . : : . : . : . . . . . . . . . . . . . . | . . . . . : . . | : . . . : . . . . . . | | . . | | .
JSR1       51 M Y L M M T V K N L R T P G N F L V L N L A L S D F G M L F F M M P T M S I N C F A E T W V I G P F

JSR1_sol  101 E C E R Y G K L G S E L G S S L I L N I N T L A K Y I E E E M K R E S D E K I P E E E I K K Q V E E
              . | | . | | . : | | . . | | : . | . : : . . : . . . . . . . : . : . . . . | . . . : . . . . . . . .
JSR1      101 M C E L Y G M I G S L F G S A S I W S L V M I T L D R Y N V I V K G M A G K P L T K V G A L L R M L

JSR1_sol  151 L E K R E K E L S S F P L T G D K G R Y V P E G S M T S C T I D Y I D T S E G P K E Y L K R Y A E E
              . . . . . . . . . : . . . . . . . . | | | | | | | | | | | | | | | | | | : . . | . . | | . . | | . .
JSR1      151 F V W I W S L G W T I A P M Y G W S R Y V P E G S M T S C T I D Y I D T A I N P M S Y L I A Y A I F

JSR1_sol  201 V Y I K P I E E A E K I V K K I K K K L E K E K K E L E K L A S L L P E G P E P L K E K K K K L E E
              | | . . | : . . . . . . . . . | . . : : . . . : | . | . : . | . . : . . . . . . . . | . . | | . . .
JSR1      201 V Y F V P L F I I I Y C Y A F I V M Q V A A H E K S L R E Q A K K M N I K S L R S N E D N K K A S A

JSR1_sol  251 I E S R V E L I E K M I K I W K E A W T P Y L K L S F D G I N S D R T N L T P E N S V K G A I K A K
              . . . . . : : . . . . | . . | . . | | | | | | . | | | . | | . | | | | . | | | . . | | . | | | . | |
JSR1      251 E F R L A K V A F M T I C C W F M A W T P Y L T L S F L G I F S D R T W L T P M T S V W G A I F A K

JSR1_sol  301 L G A I K I I E L I F E E Y A K K V A N D P S I I
              . . | . . . . . : . . . . : . | . . | . . . . . .
JSR1      301 A S A C Y N P I V Y G I S H P K Y R A A L H D K F
```

**Identity:** 94/325 (28.9%)
**Similarity:** 131/325 (40.3%)
**Gaps:** 0/325 (0.0%)

**Supplementary Figure 3: Sequence alignments for the soluble analogues against the target membrane proteins.** Performed using EMBOSS NEEDLE webserver using an EBLOSUM64 matrix.

SRII$_{sol}$

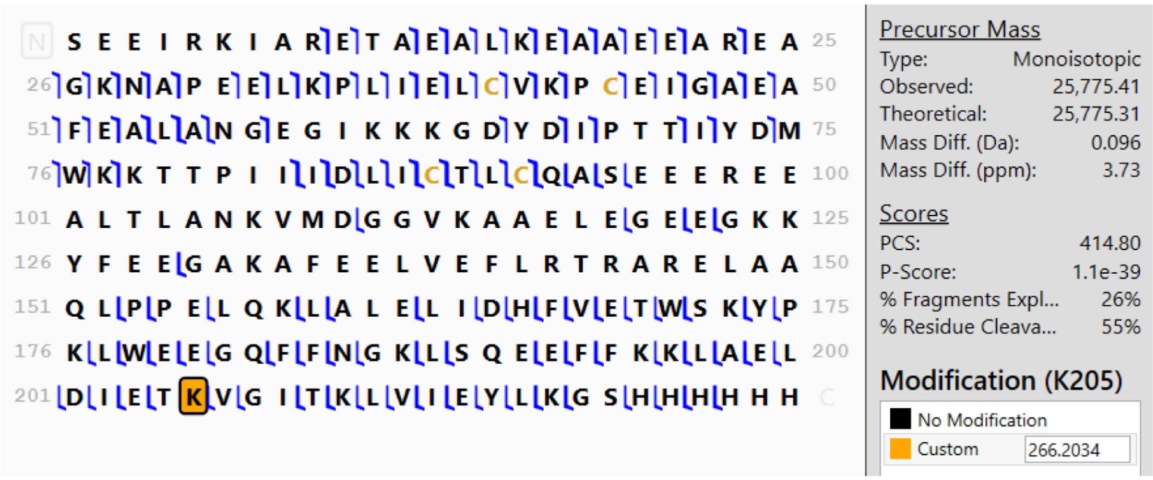


BovR$_{sol}$

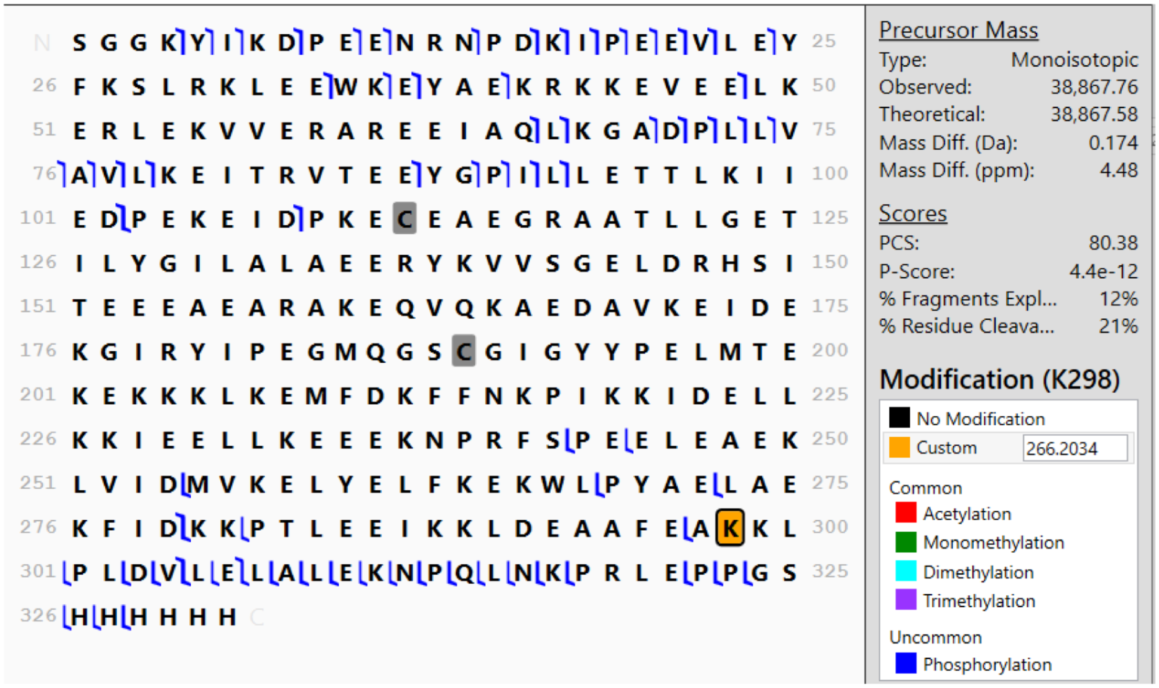


**Supplementary Figure 4: Top-down fragmentation MS data.** Detected b- and y-fragment ions are denoted in blue. Grey squares on cysteine residues indicate the presence of disulphide bond, while orange squares on lysines indicate the addition of 266.2034 Da that corresponds to retinal attachment (addition of $C_{20}H_{26}$).

| | |
|---|---|
| Wavelength | 0.9655 |
| Resolution range | 36.93 - 1.758 (1.87 - 1.76) |
| Space group | P 1 21 1 |
| Unit cell | 33.762 51.15 53.799 90 97.193 90 |
| Total reflections | 107893 |
| Unique reflections | 17443 (2346) |
| Multiplicity | 5.9 |
| Completeness (%) | 95.77 (77.53) |
| Mean I/sigma(I) | 11.4 |
| Wilson B-factor | 35.85 |
| CC1/2 | 0.999 |
| Reflections used in refinement | 17443 (2346) |
| Reflections used for R-free | 895 (103) |
| R-work | 0.1932 (0.2491) |
| R-free | 0.2211 (0.3458) |
| Number of non-hydrogen atoms | 1593 |
| macromolecules | 1558 |
| ligands | 20 |
| solvent | 15 |
| Protein residues | 195 |
| Nucleic acid bases | 0 |
| RMS(bonds) | 0.01 |
| RMS(angles) | 1.09 |
| Ramachandran favored (%) | 98.94 |
| Ramachandran allowed (%) | 1.06 |
| Ramachandran outliers (%) | 0 |
| Rotamer outliers (%) | 1.23 |
| Clashscore | 0 |
| Average B-factor | 52.56 |
| macromolecules | 52.79 |
| ligands | 42 |
| solvent | 43.23 |

**Supplementary Table 3: X-ray collection data and refinement statistics for the crystal structure of $SRII_{sol}$.**
